# Effect of tryptophan starvation on inclusion membrane composition and chlamydial-host interactions

**DOI:** 10.1101/2024.11.26.625498

**Authors:** Camille M. Riffaud-Widner, Ray E. Widner, Scot P. Ouellette, Elizabeth A. Rucks

**Affiliations:** Department of Pathology, Microbiology, and Immunology, College of Medicine, University of Nebraska Medical Center, Omaha, Nebraska, USA 68198

**Keywords:** *Chlamydia trachomatis*, tryptophan, interferon-γ, persistence, inclusion membrane, Inc, intracellular trafficking

## Abstract

*Chlamydia* is an obligate intracellular bacterial pathogen that develops within a membrane-bound vacuole called an inclusion. Throughout its developmental cycle, *Chlamydia* modifies the inclusion membrane (IM) with type III secreted (T3S) membrane proteins, known as inclusion membrane proteins (Incs). Via the IM, *Chlamydia* manipulates the host cell to acquire lipids and nutrients necessary for its growth. One key nutrient is tryptophan (Trp). As a Trp auxotroph, *Chlamydia* is very sensitive to Trp starvation and, in response to low Trp levels induced by the immune response, enters a viable but nonreplicating state called persistence. To maintain viability during persistence, *Chlamydia* must necessarily maintain both the integrity of the IM and its ability to modify host cell responses, but how Trp starvation affects IM composition and subsequent interactions with the host cell remains poorly understood. We hypothesize that, under Trp starvation conditions, Inc expression/stability or T3S function during persistence alters IM composition but that key host-*Chlamydia* interactions will be preserved. To examine host-*Chlamydia* interactions during persistence, we examined sphingomyelin, cholesterol, and transferrin trafficking to the inclusion, as well as localization of host proteins that bind to specific Incs. We identified IM composition changes during persistence by monitoring endogenous Inc abundance at the IM. Chlamydial T3S is generally functional during persistence. Specific changes in Inc composition in the IM can be linked to Trp content of a specific Inc or effector-specific defects in chlamydial T3S. Overall, our findings reveal that critical host-*Chlamydia* interactions are maintained during persistence mediated by Trp starvation.

## Introduction

The Gram-negative bacterium *C. trachomatis* from the *Chlamydiaceae* family is an obligate intracellular pathogen and the leading cause of bacterial sexually transmitted infections and preventable infectious blindness. *Chlamydia* utilizes a biphasic developmental cycle within a host cell to alternate between two distinct morphological and functional forms: a smaller, metabolically quiescent, infectious elementary body (EB), and a larger, non-infectious replicative reticulate body (RB) (1). Importantly, the RBs reside in a *Chlamydia*-containing vacuole called the inclusion, which offers a protective and stable growth niche. The inclusion membrane (IM) is modified by type III secretion (T3S) effectors that insert into the IM. These effectors are known as inclusion membrane proteins (Incs) (2). Inclusion maturation is necessary for chlamydial growth, which includes blocking fusion with lysosomes and redirecting host nutrients to the inclusion. Via the IM, *C. trachomatis* interacts with multiple subcellular compartments, such as the endoplasmic reticulum and the Golgi, to scavenge amino acids and lipids (2). The structural integrity of the IM is essential for chlamydial development (3), suggesting that an intact IM and the presence of specific Incs are necessary for chlamydial viability. Incs contain two or more hydrophobic transmembrane domains and are temporally expressed during the developmental cycle (4–7). The specific function of most Incs remains unknown, but studies have demonstrated that some Incs organize the localization of other Incs within the IM and/or recruit eukaryotic proteins to the inclusion (8–13). Some interactions between different host factors and Incs have been described. For example, IncE interacts with the host protein, sorting nexin-6 (SNX6) (10), and IncD interacts with the host ceramide transfer protein, CERT (12, 13).

*Chlamydia* has adapted to obligate intracellular dependence and acquires essential nutrients from the host cell, including amino acids. As a consequence of chlamydial evolution and reduction of its genome size, *Chlamydia* is auxotrophic for tryptophan (Trp) and most other amino acids (14), which highlights the vulnerability of *C. trachomatis* to amino acid limitation (15, 16). Within its intracellular niche, *Chlamydia* is protected from osmotic stress and the host immune system but is highly susceptible to amino acid starvation. To survive this environmental stress, *Chlamydia spp.* enter a survival state called persistence. Persistence was initially characterized for free-living bacteria exposed to antibiotics (17) but is now appreciated as a common stress response, especially for intracellular pathogenic bacteria exposed to stressors including antimicrobial host responses such as amino acid limitation or iron starvation (16). During persistence, *Chlamydia spp.* are viable, but quiescent, and can survive for a long period of time until the inducing stress is removed (18). Chlamydial persistence is induced by a mechanism that does not involve the stringent response (19–23). Further, the inclusion must remain intact during persistence to protect the quiescent *Chlamydia* from intracellular antimicrobial processes; however, the composition of the IM during persistence has yet to be characterized.

Recently, Brockett and Liechti (24) reported an association between the aberrant phenotype induced by the iron chelator, bipyridyl, or compound C1 (incorrectly referenced as a T3SS inhibitor (25)) and altered development of the inclusion. In their study, bipyridyl treatment led to the absence of IncA, and subsequent lack of inclusion fusion, and CT813/**InaC** (mid-cycle Incs) at the IM of the aberrant *Chlamydia* (24). These data indicate that chlamydial aberrance induced by metal chelation may result in changes to the IM composition and thus, chlamydial-host interactions. However, no systematic study has been performed to determine the effects of amino acid starvation on *Chlamydia*-host interactions or IM composition.

During an infection, *C. trachomatis* can encounter Trp starvation as a result of interferon-γ (IFNγ) binding to its receptor on the host cell, which activates the production of indoleamine 2,3-dioxygenase (IDO) that catabolizes Trp into *N′*-formylkynurenine (26, 27). *Chlamydia* is auxotrophic for Trp and responds to the low Trp environment by the development of a persistence phenotype (14, 16, 28). Therefore, IFNγ is commonly used to trigger persistence of *Chlamydia* in cell culture models (16, 19–22, 28–31). In *Chlamydia*, persistence is defined as a viable but non-infectious reversible state of chlamydial development that is characterized by morphologically aberrant, enlarged RBs (aRB, size of ≈2 to 10 μm), lack of cell division, and severe reduction/elimination in progeny production (18). *Chlamydia* responds to IFNγ-mediated Trp starvation by a global increase of transcription and a decrease in translation (19). Specifically, the transcription of genes containing Trp codons was generally increased in response to IFNγ treatment whereas the transcripts containing no or few Trp codons were generally unchanged (20). We further demonstrated that this may be a conserved mechanism among Trp auxotrophs in response to Trp starvation (20, 22). In *C. trachomatis* serovar L2, the proteome average of Trp is 0.94% (22). More recently, work from our laboratories demonstrated that chlamydial cell division is blocked during Trp starvation-mediated persistence by the inability to translate Trp-rich (above 1.2%) cell division proteins such as PBP2, RodA, FtsI/PBP3, and MraY. Furthermore, the Trp-poor (between 0 and 0.96%) protein, MurG, and the Trp-neutral (between 0.96% and 1.2%) protein, FtsL, were still expressed, linking Trp content of a protein of interest and its expression during low Trp conditions (23). In the same study, we demonstrated that the Trp content significantly decreased the expression of a wild-type protein containing a WW motif such as the division protein RodZ during persistence whereas expression of an isoform lacking Trp codons was maintained under the same conditions (23). Overall, these findings provide a mechanistic explanation for how *Chlamydia* blocks cell division during Trp starvation. What remains undefined is which host-*Chlamydia* interactions are preserved, if T3S is functional, and which Incs are present on the IM during persistence.

Given the large number of predicted *inc* genes for *C. trachomatis* (60 to 80, which is 7-8% of the genome) (4, 32), we wanted to understand how Inc function is affected by persistence mediated by Trp starvation. In tissue culture, we induced chlamydial persistence using two different mechanisms. Firstly, we used IFNγ, which is the most used and physiologically relevant methodology to induce Trp starvation-mediated persistence in *Chlamydia*. However, as described above, IFNγ treatment alters the environment of both the infected cell and *Chlamydia*. To better understand chlamydial-specific responses to Trp-starvation, we used indolmycin, a prokaryotic tryptophanyl-tRNA synthetase inhibitor, which acts as a Trp analog to mimic Trp depletion specifically within *Chlamydia* (33). We hypothesized that Inc expression, secretion, and abundance at the IM depend on their Trp content and that altering the composition of the IM would result in changes in host-*Chlamydia* interactions during persistence. To examine this, we initially assessed the recruitment of key lipids and trafficking proteins to the inclusion. We next examined the localization of selected endogenous Incs during normal growth and during Trp starvation conditions mediated by IFNγ or indolmycin. Our data revealed that Inc localization to the IM is not necessarily linked to Trp codon content, but other factors including T3S functionality and Inc stability also impact IM content. Overall, our data demonstrate that long-term survival of persistent *Chlamydia* is supported by the recruitment of essential host factors and specific changes in Inc composition of the IM.

## Results

### Effect of tryptophan limitation on lipid trafficking and Rab6 recruitment to the chlamydial inclusion

From within the inclusion, *Chlamydia* intercepts Golgi-derived exocytic vesicles that contain sphingomyelin (SM) (34–36). Previous studies have found that SM is required for chlamydial growth (37), but it has not been shown whether persistent *Chlamydia* recruit SM and other Golgi-related cargo to the inclusion. Golgi-derived SM trafficking to the inclusion is determined by incubating infected cells with NBD-ceramide (38). In the Golgi, NBD-ceramide is metabolized into NBD-glucosylceramide and NBD-sphingomyelin (NBD-SM), which both traffic to the plasma membrane (39, 40). NBD-lipids can be removed from the plasma membrane through a process known as back-exchange, which is achieved with medium containing FBS or BSA that binds to the NBD-moiety and irreversibly strips the lipid from the plasma membrane. NBD-SM incorporation into the inclusion and chlamydial organisms is recorded via live-cell imaging (Fig. 1A). For these studies, HEp-2 cells were either pre-treated or not with IFNγ, then infected with *C. trachomatis* serovar L2 that constitutively expressed mCherry (41). At 10 hpi, monolayers remained untreated or treated with IFNγ or indolmycin, and examination of chlamydial recruitment of NBD-SM followed the indicated timelines (Fig. S1A). Additionally, untreated monolayers were processed at 12 hpi for live-cell imaging of inclusion retention of SM, as inclusions at these timepoints are similar in size to those formed by persistent organisms at 24 hpi (Fig. S1B). Trp-starvation induced by IFNγ or indolmycin results in few chlamydial organisms and smaller inclusions, and, since NBD-SM photobleaches relatively quickly, mCherry signal was used to focus on inclusions. Images were analyzed in ImageJ to determine the integrated density of NBD-SM, which was then normalized to inclusion size (Fig. 1B). Inclusions containing persistent *Chlamydia* were significantly brighter (greater intensity of NBD-SM) than the inclusions of the untreated control at 12 hpi, similar in brightness to the untreated inclusions at 27 hpi (p<0.0001) (Fig. 1B). To our knowledge, these are the first studies to demonstrate that *Chlamydia* acquires SM during persistence.

**Figure 1.**
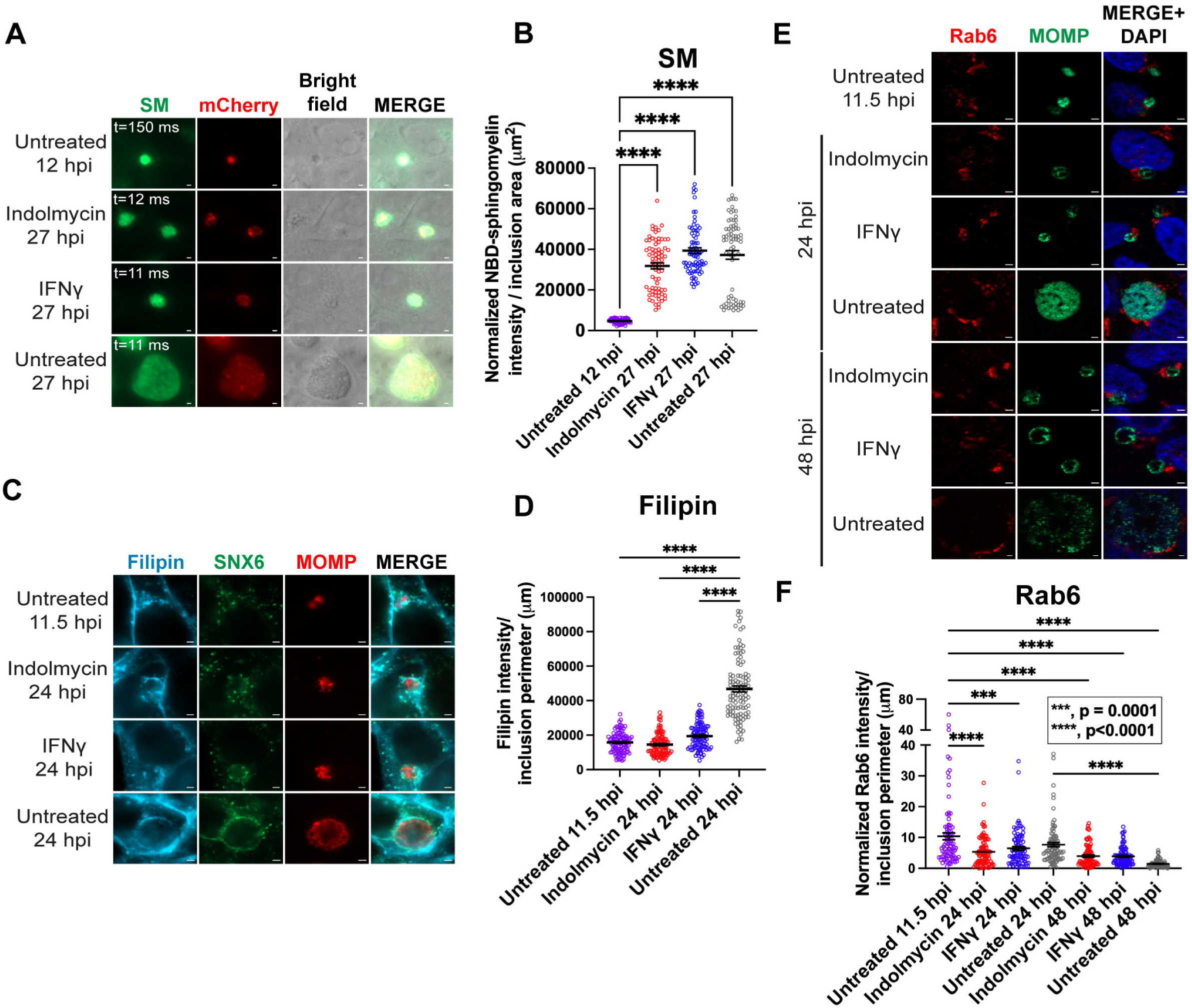
Golgi trafficking to the inclusion is maintained during Trp starvation. **(A)** HEp-2 cells were infected with *C. trachomatis* serovar L2 pBOMBmC-GFP(VAA) and treated or not for aberrancy. Cells were labeled with NBD-ceramide as described in Materials and Methods. Live-cell images were captured in the brightfield, 488, and 594 fluorescent channels at 63x; scale bar = 2 μm. **(B)** Chlamydial acquisition of NBD-sphingomyelin (NBD-SM) was quantified as described in Materials and Methods. Individual values representing the intensity of NBD-SM normalized to exposure time and divided by the inclusion area (μm^2^) with mean and standard error of the mean (SEM) were graphed and analyzed by an ordinary one-way ANOVA with Šidák’s multiple comparisons test. ****, p<0.0001. Data shown are combined from 3 independent experiments (75 inclusions total per condition). **(C-F)** HEp-2 cells were infected with *C. trachomatis* serovar L2 and treated or not for Trp starvation. **(C-D)** After fixation at 11.5 hpi and 24 hpi, HEp-2 cells were incubated with filipin to detect cholesterol. Samples were imaged with a consistent exposure time across conditions and at 100x; scale bar = 2 μm. In **D**, the raw integrated density of filipin and the inclusion perimeter were determined. Raw integrated densities were divided by the inclusion perimeters, and values were graphed with mean and SEM as described above. ****, p<0.0001. Data shown are combined from 3 independent experiments (105-107 inclusions total per condition). **(E)** Samples were fixed at 11.5 hpi, 24 hpi, and 48 hpi, and processed for indirect immunofluorescence to detect Rab6 (red), MOMP (green), and DNA (blue). Scale bar = 2 μm. **(F)** Immunofluorescence images were quantified using Fiji/ImageJ software to determine the raw integrated density of Rab6, normalized to exposure time, and divided by inclusion perimeter (µm). Results were graphed and statistically analyzed using GraphPad Prism as described in B. ***, p<0.001; ****, p<0.0001. Note the break in the y-axis. Data shown are combined from 3 independent experiments (90 inclusions total per condition).

We also tested whether *Chlamydia* acquires cholesterol, which traffics from the Golgi to the chlamydial inclusion (36). As above, HEp-2 cells were infected with wild-type *C. trachomatis* L2 and treated for persistence or not and fixed at 24 hpi, then labeled with filipin to detect host cholesterol (Fig. 1C and Fig. S1C). Untreated infected monolayers were also fixed at 11.5 hpi, as these inclusions are similar in size to inclusions at 24 hpi during Trp-starvation/persistence (Fig. S1D). Consistent with induction of persistence by indolmycin or IFNγ, aRBs were apparent within the small inclusions (Fig. 1C). Compared to the untreated inclusions at 24 hpi, inclusions from indolmycin- or IFNγ-mediated persistence conditions showed significantly less filipin staining at the IM (p<0.0001), but similar filipin levels as untreated inclusions at 11.5 hpi (Fig. 1D). Thus, *C. trachomatis* still traffics cholesterol to the IM under persistence conditions.

Rab6 is a Golgi-associated low molecular weight GTPase that localizes to *C. trachomatis* serovar L2 inclusions (42) and has been demonstrated to participate in SM trafficking to the inclusion (43). To determine if Rab6 localizes to inclusions containing persistent organisms, HEp-2 cells were seeded, infected, and treated or not to induce persistence via Trp starvation as described above. Treatment of infected monolayers with either indolmycin or IFNγ and fixation at 24hpi resulted in inclusions of similar size to untreated monolayers at 11.5 hpi that contained enlarged organisms, which is consistent with induction of persistence (Fig. 1E). In all infected monolayers, Rab6 was observed in Golgi-like structures that fragmented around the inclusion (Fig. 1E), consistent with previous observations (44, 45). The intensity/integrated density of Rab6 at the inclusion was normalized to exposure times and then, to account for differences in inclusion sizes, divided by the perimeter of the inclusion. In untreated cultures, the intensity of Rab 6 is highest during early development (11.5 hpi) and lessens in intensity at later timepoints in chlamydial development (p=0.0323 compared to untreated at 24 hpi and p<0.0001 at 48 hpi). Indolmycin- and IFNγ-induced persistence conditions resulted in similar levels of Rab6 intensity at the inclusion as 24 hpi untreated cultures but was significantly less than what was observed in untreated cultures at 11.5 hpi (p<0.0001 for indolmycin and p=0.0001 for IFNγ) (Fig. 1F). To determine if Rab6 localization to inclusions in cells treated with indolmycin or IFNγ was simply delayed, persistent cultures were incubated for an additional 24 hours and fixed at 48 hpi. At 48 hpi, we observed that Rab6 was localized in a compact Golgi structure adjacent to inclusions in both indolmycin- and IFNγ-treated cells (Fig. 1E). At 48 hpi, the relative intensity of Rab6 at the IM was decreased during both IFNγ- and indolmycin-induced persistence compared to persistent monolayers fixed at 24 hpi, but not significantly (p=0.0556 for IFNγ at 24 hpi versus 48 hpi and p=0.8863 for indolmycin at 24 hpi versus 48 hpi). The overall association of Rab6 with inclusions of persistent organisms was indistinguishable between indolmycin- and IFNγ-treated cultures at either 24 or 48 hpi. Altogether, these data suggest that *C. trachomatis* can recruit the host Golgi-derived lipids, sphingomyelin and cholesterol, and the host Golgi-associated Rab GTPase, Rab6, to the inclusion during persistence, regardless of the mechanism of inducing persistence.

### Effect of tryptophan limitation on Rab11-FIP1 and transferrin trafficking to the inclusion

The inclusion also interacts with components of the endosomal recycling pathway such as Rab11-FIP1 (46) and transferrin (Tf) via the slow Tf recycling pathway (47). Rab11-FIP1 is involved in Tf trafficking via its function in coupling Rab11 and Rab4 vesicles in the slow-recycling pathway of Tf (47–49). To determine if Trp starvation induced by indolmycin or IFNγ affected transport of these markers to the inclusion, *C. trachomatis* serovar L2 was treated or not for Trp-induced persistence at 10 hpi and the localization of Rab11-FIP1 or Tf at the IM was evaluated by indirect immunofluorescence at 11.5 hpi (Rab11-FIP1 and Tf) (similarly sized inclusions to those containing persistent bacteria), 24 hpi (Rab11-FIP1 and Tf), and 48 hpi (Rab11-FIP1 only) (Fig. 2). Both Rab11-FIP1 and Tf associated with the inclusions throughout persistence, as the signal was still evident in persistent cultures at 24 hpi (Tf) and 48 hpi, (Rab11-FIP1) which are timepoints when localization of either host component wanes in untreated controls. Comparing untreated 11.5 hpi inclusions to those under persistence conditions at 24 hpi, there was significantly less Rab11-FIP1 at the inclusion under indolmycin treatment (p<0.0001) but not IFNγ treatment (p=0.2515) (Fig. 2C). At 24 hpi, there were no statistically significant differences in Rab11-FIP1 inclusion localization among untreated and persistence conditions; however, a greater intensity of Rab11-FIP1 was measured at inclusions when persistence was induced with IFNγ compared to indolmycin, but this was not statistically significant (p=0.0674). At 48 hpi, Rab11-FIP1 localization to inclusions in the untreated condition was significantly lower than both persistence conditions (p=0.0028 for indolmycin and p=0.0004 for IFNγ). Like the localization patterns of Rab11-FIP1, transferrin trafficking to the inclusion decreases over chlamydial development, comparing the untreated 11.5hpi and 24 hpi conditions (Fig. 2D, p<0.0001). Transferrin localization to the inclusions containing persistent organisms at 24 hpi was similar to what was observed in inclusions of untreated monolayers at 11.5 hpi but elevated relative to the untreated condition at 24 hpi (Fig. 2D, p=0.0081 for indolmycin and p<0.0001 for IFNγ). Overall, these data demonstrate that chlamydial interactions with the recycling endosomal pathway are maintained during persistence induced with either indolmycin or IFNγ.

**Figure 2.**
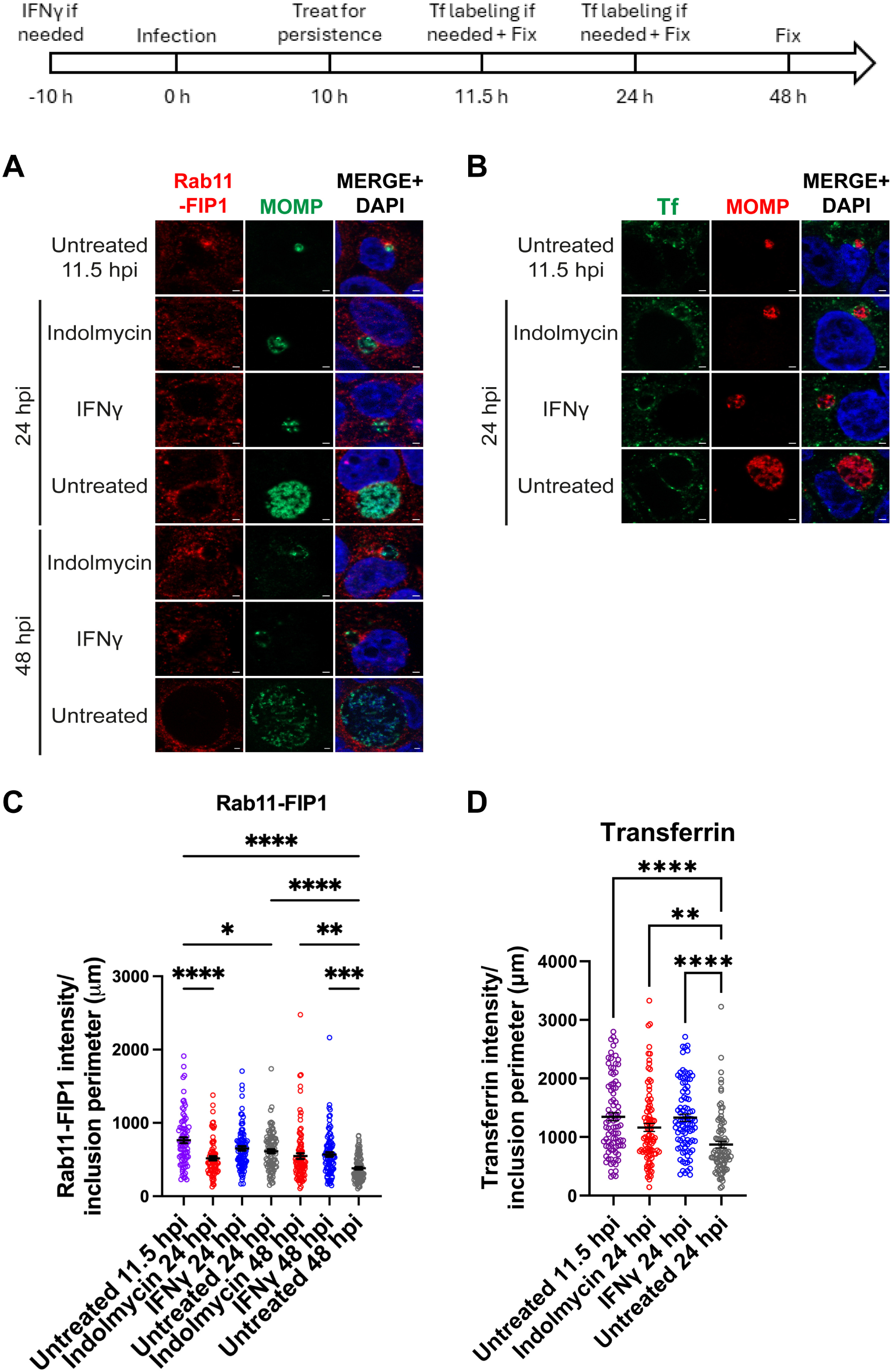
Trafficking from the recycling endosomal pathway to the chlamydial inclusion is maintained during Trp starvation. A diagram representing the experimental design and highlighting important timepoints is shown. HEp-2 cells were infected with *C. trachomatis* serovar L2 and treated or not for Trp starvation. **(A)** Samples were fixed at 11.5 hpi, 24 hpi, and 48 hpi, and processed for indirect immunofluorescence to detect Rab11-FIP1 (red), MOMP (green), and DNA (blue). Scale bar = 2 μm. **(B)** Infected cells were treated with 25 μg/mL transferrin-488 (Tf, green), as described in Methods. Cells were fixed and processed to detect MOMP (red) and DNA (blue). Scale bar = 2 μm. **(C-D)** Immunofluorescence images were quantified using Fiji/ImageJ software to determine the raw integrated density of Rab11-FIP1 (C) or transferrin-488 (D) per inclusion perimeter (µm), and results (mean and SEM) were graphed and statistically analyzed (via ordinary one-way ANOVA with Šidák’s multiple comparisons test) using GraphPad Prism. *, p<0.05; **, p<0.01; ***, p<0.001; ****, p<0.0001. Data shown are combined from 3 independent experiments (77-92 inclusions total per condition for Rab11-FIP1 and 90 inclusions total per condition for transferrin).

### Trp content of the *C. trachomatis* L2 Inc proteins included in our study

Given our previous findings that the Trp content of a protein affects its expression during Trp starvation in *Chlamydia* (20, 23), we hypothesized that the expression of Incs is similarly changed during Trp starvation as a function of their Trp content. Because antibodies against endogenous Incs are not widely available, we prioritized Incs based on the availability of reagents on-hand. Fortunately, these reagents allowed the examination of 7 Incs. The genes encoding these Incs differ in whether they are organized or not in an operon, contain Trp codons, the percentage of Trp, the presence of a tandem Trp motif (i.e., WW), and the temporal expression during the developmental cycle (Table 1). Specifically, the genes *incD*, *incE*, *incF*, and *incG* are encoded in an operon and are expressed early in the developmental cycle (3 hpi with peak of expression at 8 hpi) (7, 50). Genes encoding CT223, InaC, and IncA are not found organized in operons and represent an early Inc (CT223 (51, 52)) or mid-cycle Incs with peak of expression at 16-20 hpi (InaC and IncA) (7). Three of the Incs examined lack Trp: IncA, IncE, and IncG, while InaC has a higher percentage of Trp (2.27%) than the *C. trachomatis* L2 proteome average (0.94%), and its sequence is further enriched in Trp by the presence of a WW motif (Table 1). With the exception of InaC, the percentage of Trp in our selection of Incs is poor or neutral. Therefore, if our hypothesis is correct, then CT223, IncA, IncD, IncE, IncF, and IncG should be expressed during Trp starvation, while expression of InaC should be decreased.

**Table 1:** Tryptophan content of a selection of Inc proteins in *Chlamydia trachomatis* L2.

| Common protein name | *Chlamydia trachomatis* L2/434/Bu | Presence in an operon* | Tryptophan codons | Tryptophan motif | Percentage of Tryptophans | Expression during the developmental cycle** |
| --- | --- | --- | --- | --- | --- | --- |
| IncD | CTL0370 | yes, start of the operon | 1 | no | 0.68% | early |
| IncE | CTL0371 | yes, middle of the operon | 0 | no | 0% | early |
| IncF | CTL0372 | yes, middle of the operon | 1 | no | 0.96% | early |
| IncG | CTL0373 | yes, end of the operon | 0 | no | 0% | early |
| CT223/IpaM | CTL0476 | no | 1 | no | 0.37% | early |
| InaC | CTL0184 | no | 6 | **WW** | 2.27% | mid |
| IncA | CTL0374 | no | 0 | no | 0% | mid |

\* Based on genomic context
\*\* Times are relevant for *C. trachomatis* serovar L2 (references 6, 7, and 52)
Tryptophan motif is indicated in bold

### Effect of tryptophan limitation on localization of IncD and IncE and their binding partners, CERT and SNX6, respectively, at the inclusion membrane during Trp starvation

To determine if Trp starvation affects IncD (1 Trp) and IncE (0 Trp) protein expression on the IM, HEp-2 cells infected with wild-type *C. trachomatis* L2 were starved or not for Trp at 10 hpi with indolmycin or IFNγ, essentially as above. Consistent with indolmycin- or IFNγ-induced persistence in *Chlamydia*, we observed smaller inclusions containing aberrant organisms, which are similar in size to 11.5 hpi inclusions (Fig. 3A, C, E, and G). Subcellular localization of endogenous IncD, evaluated by indirect immunofluorescence and quantified in Image J, demonstrated that IncD levels increase over the course of chlamydial development, comparing untreated infected monolayers at 11.5 hpi to 24 hpi (Fig 3B; p<0.0001). Further, there is a similar intensity of IncD in indolmycin-induced persistent inclusions at 24 hpi compared to 11.5 hpi untreated inclusions and a modest but statistically significant increase on inclusions in IFNγ-treated cells at 24 hpi compared to 11.5 hpi untreated inclusions (p=0.012) (Fig. 3B). However, intensity of IncD in the IMs within monolayers treated with indolmycin or IFNγ was significantly less than what was observed at 24 hpi in untreated cultures (Fig. 3B; p<0.0001). Similarly to IncD, the intensity of IncE on the IM increases over the course of chlamydial development from 11.5 hpi to 24 hpi in untreated infected cultures (Fig. 3D; p<0.0007). In contrast to IncD, the intensity of IncE on the IM was similar between 24 hpi untreated inclusions and inclusions under indolmycin or IFNγ conditions. Next, known IncD and IncE binding partners, CERT (12, 13) and sorting nexin-6 (SNX6) (10), respectively, were examined for localization to the inclusion during persistence. Consistent with the expression of IncD, CERT intensity at the inclusion did not significantly change in persistence conditions compared to the untreated 11.5 hpi inclusions (Fig. 3F). Surprisingly, CERT intensity at the untreated 24 hpi inclusions was lower than for any other sample despite IncD levels on the IM being highest for the untreated 24 hpi inclusions. In contrast, SNX6 intensity at the inclusion remained relatively constant over the course of chlamydial development with a modest, but not statistically significant, increase of intensity at 24 hpi compared to 11.5 hpi in untreated infected monolayers (Fig. 3H). Further, SNX6 intensity at the inclusion during persistence induced with indolmycin or IFNγ was similar to that of normal growth conditions. These data demonstrate that during persistence, chlamydial T3S is still functional and that Inc-eukaryotic protein relationships are maintained. These findings are also consistent with continued expression of proteins with little or no Trp; however, the expression of both *incD* and *incE* would have commenced before the onset of persistence. Therefore, the link between Inc expression on the IM during persistence and their Trp content is not resolved by these data.

**Figure 3.**
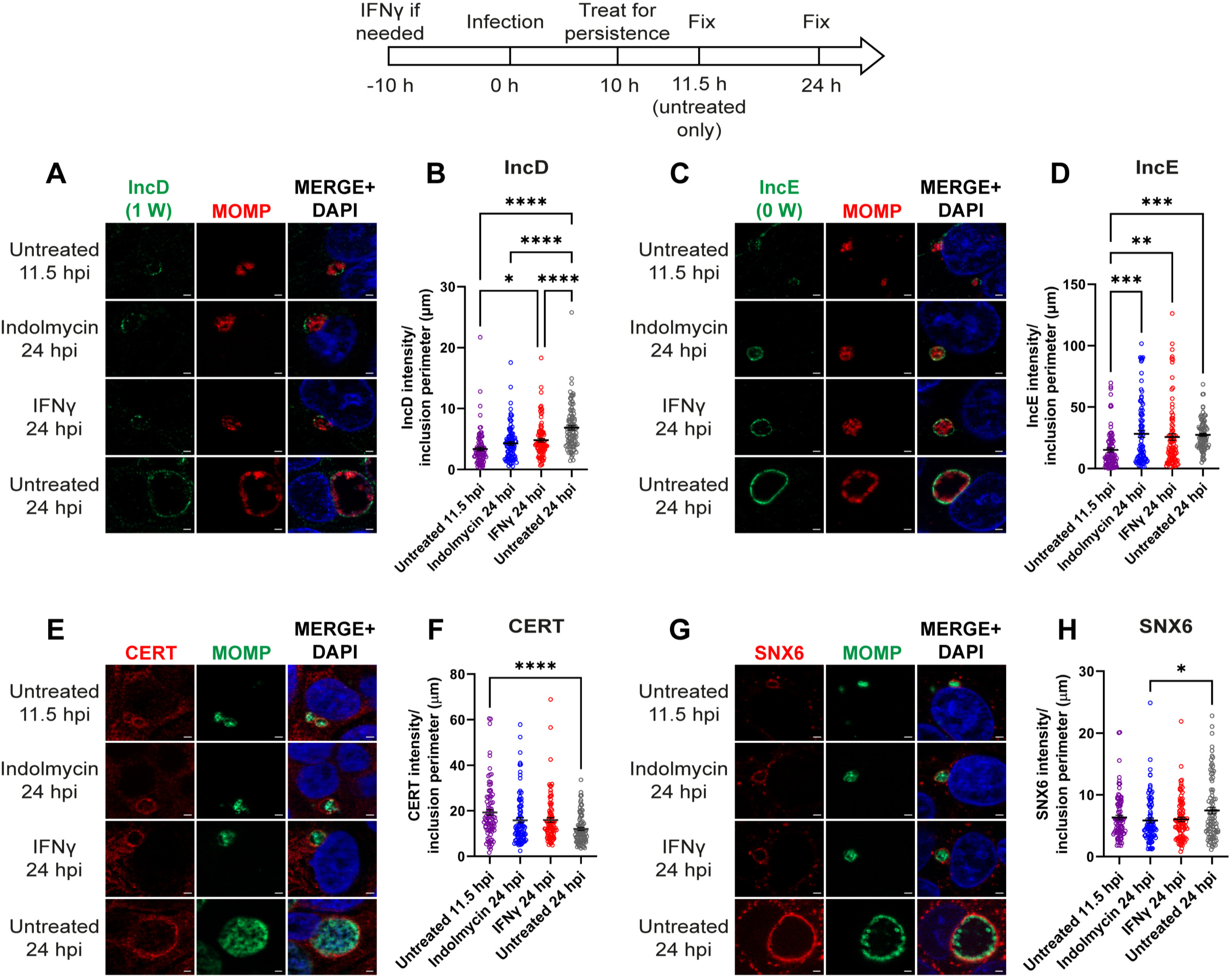
IncD and IncE localize to the inclusion membrane during Trp starvation. A diagram representing the experimental design and highlighting important timepoints is shown. HEp-2 cells were infected with *C. trachomatis* serovar L2 treated or not with indolmycin in ICM or IFNγ to induce aberrancy. **(A,** C) Samples were fixed at 11.5 hpi and 24 hpi and processed for indirect immunofluorescence to detect IncD or IncE (green), MOMP (red), and DNA (blue). (E**, G)** Samples were fixed at 11.5 hpi and 24 hpi and processed for indirect immunofluorescence to detect CERT or SNX6 (red), MOMP (green), and DNA (blue). Scale bar = 2 μm. **(B, D, F, and H)** Immunofluorescence images were quantified using Fiji/ImageJ software to determine the raw integrated density of IncD, IncE, CERT, or SNX6, normalized to exposure time, and divided by inclusion perimeter (µm). Results (mean and SEM) were graphed and statistically analyzed (ordinary one-way ANOVA with Šidák’s multiple comparisons test) using GraphPad Prism. *, p<0.05; **, p<0.01; ***, p<0.001; ****, p<0.0001. Data shown are combined from 3 independent experiments (90 inclusions total per condition).

### Effect of tryptophan limitation on inclusion membrane localization of endogenous Incs: IncG, CT223, InaC, and IncA

To better examine how temporal expression might impact Inc expression on the IM, additional early Incs (IncG (0 Trp) and CT223 (1 Trp)) and mid-cycle Incs (InaC (6 Trp) and IncA (0 Trp)) were examined in HEp-2 cells infected with *C. trachomatis* serovar L2 and starved or not for Trp (as described above and depicted in Fig. 4). Infected cells were fixed at the indicated timepoints and the intensity of Inc fluorescence at the IM was quantified and graphed from each condition. To account for differences in inclusion size, intensity measurements were divided by the inclusion perimeter (Fig. 4B). Consistent with known temporal expression patterns, the early Incs, IncG and CT223, are expressed on the IM at 11.5 hpi while mid-cycle Incs, InaC and IncA, are undetectable at the IM at 11.5 hpi. Therefore, we used the peak expression timepoint of each of these Incs for the untreated control condition in our analysis: 11.5 hpi for IncG and CT223, 16 hpi for InaC, and 24 hpi for IncA (Fig. 4A). In *Chlamydia* cultured in HEp-2 cells under normal growth conditions, CT223, IncG, and InaC are all detected at 24 hpi (Fig. S2). During persistence, IncG intensity was similar between untreated 11.5 hpi and indolmycin-induced persistant inclusions, but significantly increased during IFNγ treatment (p=0.0052) (Fig. 4B). Intensity of CT223 in the IM was similar across all conditions measured, suggesting that the amount of CT223 that is already inserted at the time of induction of persistence remains constant. In contrast, InaC (6 Trp) and IncA (0 Trp) intensities were significantly reduced at the IM during persistence compared to the untreated 16 hpi and untreated 24 hpi conditions, respectively (p<0.0001) (Fig. 4B). Overall, these data suggest that other factors, independent of Trp content, may influence expression and/or T3S of Inc proteins. Specifically, the expression of IncG (0 Trp), CT223 (1 Trp), and InaC (6 Trp) correlate with their Trp content, but it was surprising that IncA (0 Trp) was absent from the IM during persistence. These data suggest that expression of *inc* genes may in part be a function of Trp codon content, but additional factors related to T3S functionality, more generally, may also play a role.

**Figure 4.**
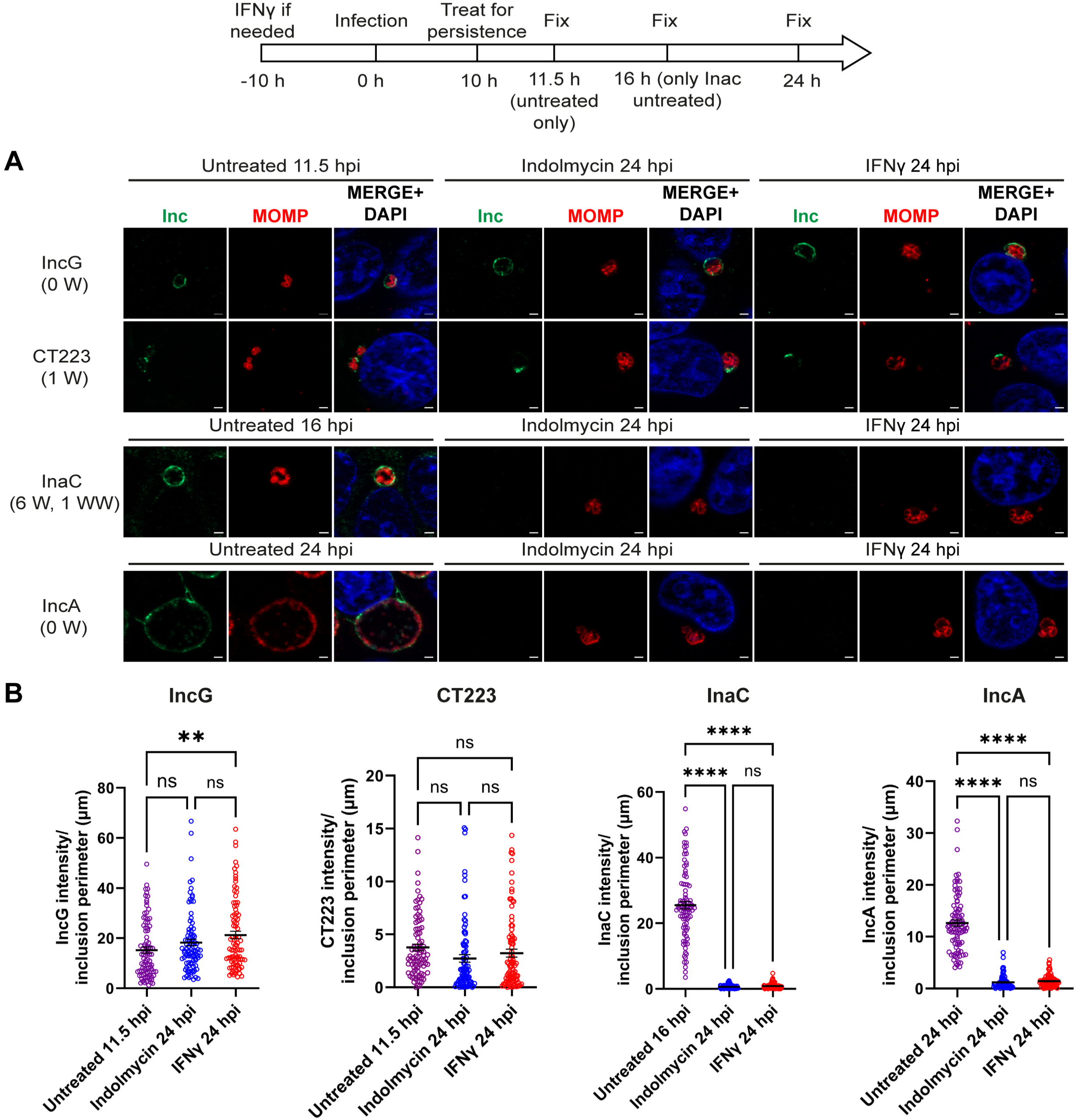
Temporal expression impacts Inc localization to the IM during Trp starvation. A diagram representing the experimental design and highlighting important timepoints is shown. HEp-2 cells were infected with *C. trachomatis* serovar L2 and treated or not for Trp starvation mediated by indolmycin in ICM or IFNγ. **(A)** Samples were fixed at 11.5 hpi, 16 hpi, and 24 hpi and processed for indirect immunofluorescence to detect Incs (green), MOMP (red), and DNA (blue). Scale bar = 2 μm. **(B)** Immunofluorescence images were quantified using Fiji/ImageJ software to determine the raw integrated density of Incs, normalized to exposure time, and divided by inclusion perimeter (µm), and results (mean and SEM) were graphed and statistically analyzed (ordinary one-way ANOVA with Šidák’s multiple comparisons test) using GraphPad Prism. ns, non-significant; **, p<0.01; ****, p<0.0001. Data shown are combined from 3 independent experiments (90 inclusions total per condition).

### Effect of tryptophan limitation on Inc secretion during persistence

The chlamydial T3S apparatus is constructed by about 26 individual structural proteins and 11 possible chaperone proteins, all of which are temporally expressed (Suppl. Table S1) (recently reviewed in (53)). The functionality of chlamydial T3S has not been tested during chlamydial persistence induced with indolmycin or IFNγ. Therefore, we designed experiments leveraging chlamydial transformants that carried plasmids for inducible expression of Incs such that we could control the expression of a given Inc during persistence. HEp-2 cells were infected with *C. trachomatis* strains carrying inducible IncA-FLAG, IncF-FLAG, or IncG-FLAG and were starved or not for Trp at 10 hpi. At 24 hpi, aTc was added to the medium to induce expression of the Inc-FLAG constructs to test secretion, then cells were fixed at 30 hpi and processed for indirect immunofluorescence (Fig. 5). Untreated, infected cells were fixed at 10 hpi to account for any leaky expression (Fig. 5A, top row). T3S of Inc-FLAG constructs was determined by subcellular localization of the construct to the IM, as in 30 hpi control images (Fig. 5A, second row). Chlamydial persistence was confirmed by the presence of enlarged organisms within small inclusions of monolayers treated with indolmycin or IFNγ (Fig. 5A and Fig. S3). Based on integrated density measurements, the Inc-FLAG constructs demonstrated limited to no leaky expression at 10 hpi (Fig. 5B). Induction of IncA-FLAG expression during persistence resulted in detection of IncA-FLAG at the IM at levels greater than the 10 hpi control (p<0.0001) but less than the 30 hpi control (p<0.0001) (Fig. 5A and B). However, neither IncF-FLAG nor IncG-FLAG was visible within organisms or on the IM (Fig. 5A), nor were they quantifiable on the IM compared to the 30 hpi untreated control (for IncF-FLAG, p=0.0007 for indolmycin and p=0.0004 for IFNγ; for IncG-FLAG, p<0.0001 for both persistence conditions) (Fig. 5B). These data indicate that the Inc composition of the IM during chlamydial persistence may be impacted by both the Trp content of the individual Inc, Inc stability within the organisms and/or on the IM, and/or the availability of components required for T3S or expression of specific effectors.

**Figure 5.**
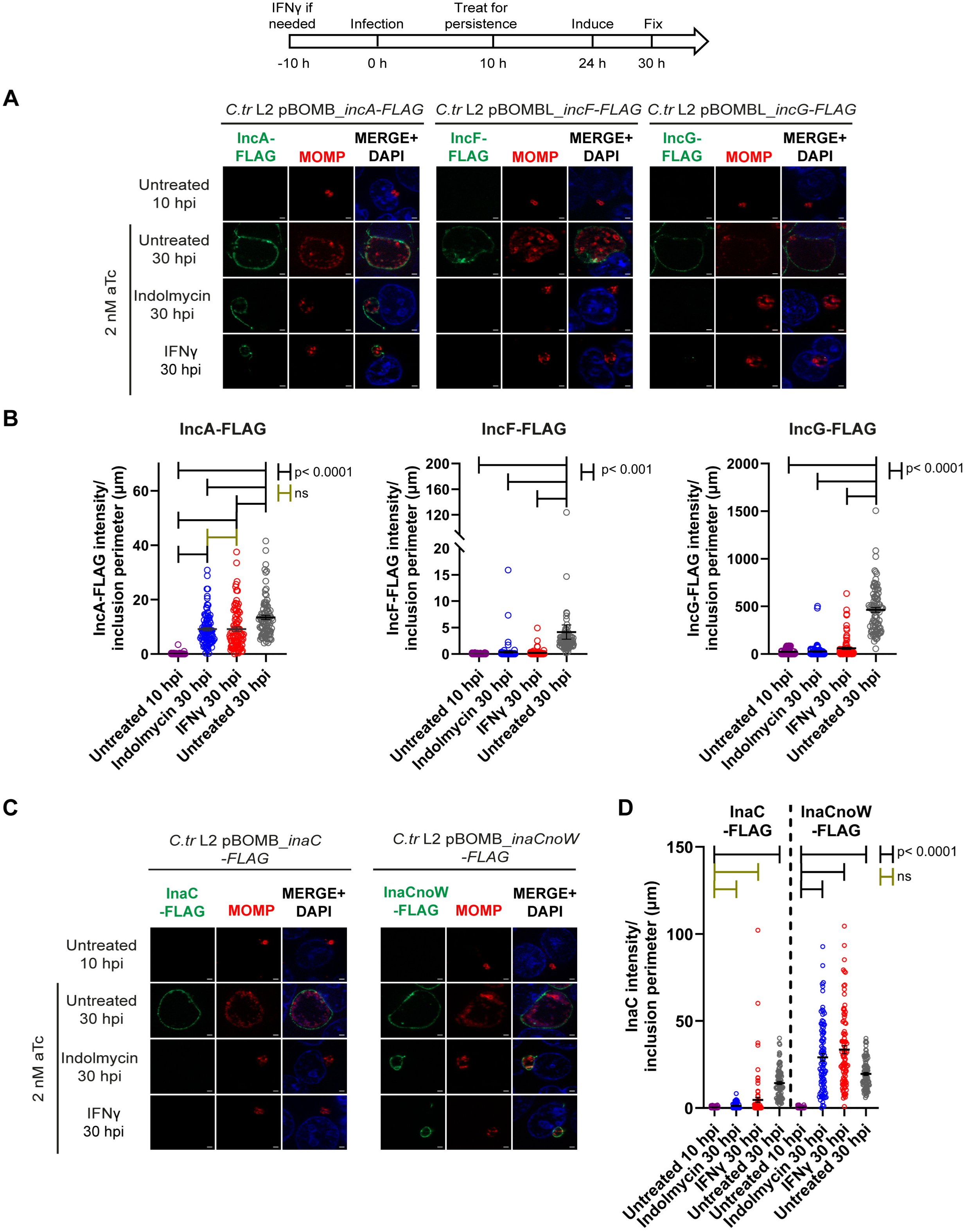
Type III secretion of Incs is differentially inhibited during Trp starvation. A diagram representing the experimental design and highlighting important timepoints is shown. At 24 hpi, 2 nM aTc was added to induce construct expression. **(A, C)** Samples were fixed at 10 hpi or 30 hpi and processed for indirect immunofluorescence to detect FLAG (green), MOMP (red), and DNA (blue). Scale bar = 2 μm. **(B, D)** Immunofluorescent images were quantified using Fiji/ImageJ software to determine the raw integrated density of FLAG tagged protein, normalized to exposure time if necessary (IncA-FLAG, IncF-FLAG, InaC-FLAG, and InaCnoW-FLAG), and divided by inclusion perimeter (µm), and results (mean and SEM) were graphed and statistically analyzed (ordinary one-way ANOVA with Šidák’s multiple comparisons test) using GraphPad Prism. Gold brackets denote ns, non-significant. Black brackets indicate statistical significance: p<0.0001 for IncA-FLAG, IncG-FLAG, InaC-FLAG, and InaCnoW-FLAG, or p<0.001 for IncF-FLAG. Note the break in the y-axis for IncF-FLAG. Data shown are combined from 3 independent experiments (90 inclusions total per condition).

Given its Trp codon content, including a WW motif, InaC is an Inc highly enriched for Trp codons. We sought to determine whether Trp starvation effectively blocked InaC-FLAG detection at the IM due to decreased expression or a defect in T3S. Therefore, we compared the IM localization of wild-type InaC-FLAG protein to an InaCnoW-FLAG mutant, wherein each Trp codon was substituted with a phenylalanine (Phe, F) codon. HEp2 cells were infected with *C. trachomatis* L2 inducible wild-type InaC-FLAG or mutant InaCnoW-FLAG transformants, then treated or not for persistence as described above. At 24 hpi, aTc was added to induce expression of InaC-FLAG or InaCnoW-FLAG, then cells were fixed at 30 hpi. Monolayers were imaged (Fig. 5C) and Inc-FLAG construct intensity at the IM was measured in Image J and divided by inclusion perimeter (Fig. 5D). Consistent with the results indicating that endogenous InaC is not expressed on the IM during persistence, InaC-FLAG is also not detected on the IM during persistence induced with indolmycin or IFNγ (Fig. 5C and D). Compared to inclusions from the untreated 30 hpi sample, inclusions in persistence conditions had significantly reduced levels of InaC-FLAG on the IM (p<0.0001). However, InaCnoW-FLAG was detected on the IM during persistence at levels significantly increased compared to the untreated 10 hpi control (Fig. 5C and D). In fact, inclusions from both persistence conditions had significantly increased InaCnoW-FLAG on the IM compared to inclusions from the untreated 30 hpi condition (p<0.0001). Finally, there was significantly more InaCnoW-FLAG on the IM during indolmycin and IFNγ treatments than there was wild-type InaC-FLAG during those same persistence conditions (p<0.0001). These data indicate that the T3S components required for InaC secretion are present during persistence, but the Trp content of InaC dictates whether it is expressed during Trp starvation.

### Effect of tryptophan limitation on Inc stability at the inclusion membrane during persistence

To determine if chlamydial persistence caused by Trp starvation mediated by indolmycin or IFNγ impacted the stability of Incs at the IM, we assessed the detection of Inc-FLAG constructs at the IM using pulse-chase conditions. To assess this parameter, HEp-2 cells were infected with *C. trachomatis* strains carrying inducible IncA-FLAG, IncF-FLAG, or IncG-FLAG, and construct expression was induced with addition of aTc at 7 hpi. At 10 hpi, the infected cells were starved or not for Trp, and aTc was also removed from the culture medium. Samples were fixed at 10 hpi to determine the amount of Inc-FLAG on the IM at the onset of persistence or 14 hours later to determine the amount of Inc-FLAG on the IM after aTc removal and during Trp starvation (Fig. 6). Fixed samples were processed for indirect immunofluorescence (Fig. 6A), and images were quantified as described above (Fig. 6B). Induction of chlamydial persistence was confirmed by the presence of enlarged bacteria (Fig. 6A) within quantifiably small inclusions (Fig. S4). During Trp-starvation induced persistence, IncA-FLAG was evident on the IM during indolmycin-induced persistence but was reduced during IFNγ-mediated persistence, while levels of IncF-FLAG and IncG-FLAG were reduced compared to untreated cells fixed at 10 hpi (Fig. 6A and B). Comparison of Inc-FLAG expression during persistence induced with indolmycin versus IFNγ compared to the 10 hpi controls showed that IncA-FLAG expression was significantly reduced only during IFNγ-induced persistence (p=0.0001), while levels of IncA-FLAG during indolmycin-induced persistence were similar to the 10 hpi control (Fig. 6B). IncA-FLAG stability was also significantly decreased in IFNγ-induced persistence when compared to indolmycin-induced persistence (p<0.0001). Similarly, less IncF-FLAG and IncG-FLAG levels were detected on the IM during IFNγ-induced persistence compared to indolmycin-induced persistence (Fig. 6B). IncF-FLAG levels were significantly reduced under both persistence conditions compared to the untreated 10 hpi control (p<0.0001), and the IFNγ treatment showed a significant decrease compared to the indolmycin treatment (p=0.0027). IncG-FLAG also showed decreased stability during persistence, with a more modest, statistically insignificant decrease during indolmycin treatment (p=0.1236) and a greater, statistically significant decrease during IFNγ treatment (p=0.0037). Unlike IncA-FLAG and IncF-FLAG, there was no statistically significant difference between persistence conditions for IncG-FLAG. These data suggest that host processes associated with IFNγ treatment may impact Inc stability on the IM.

**Figure 6.**
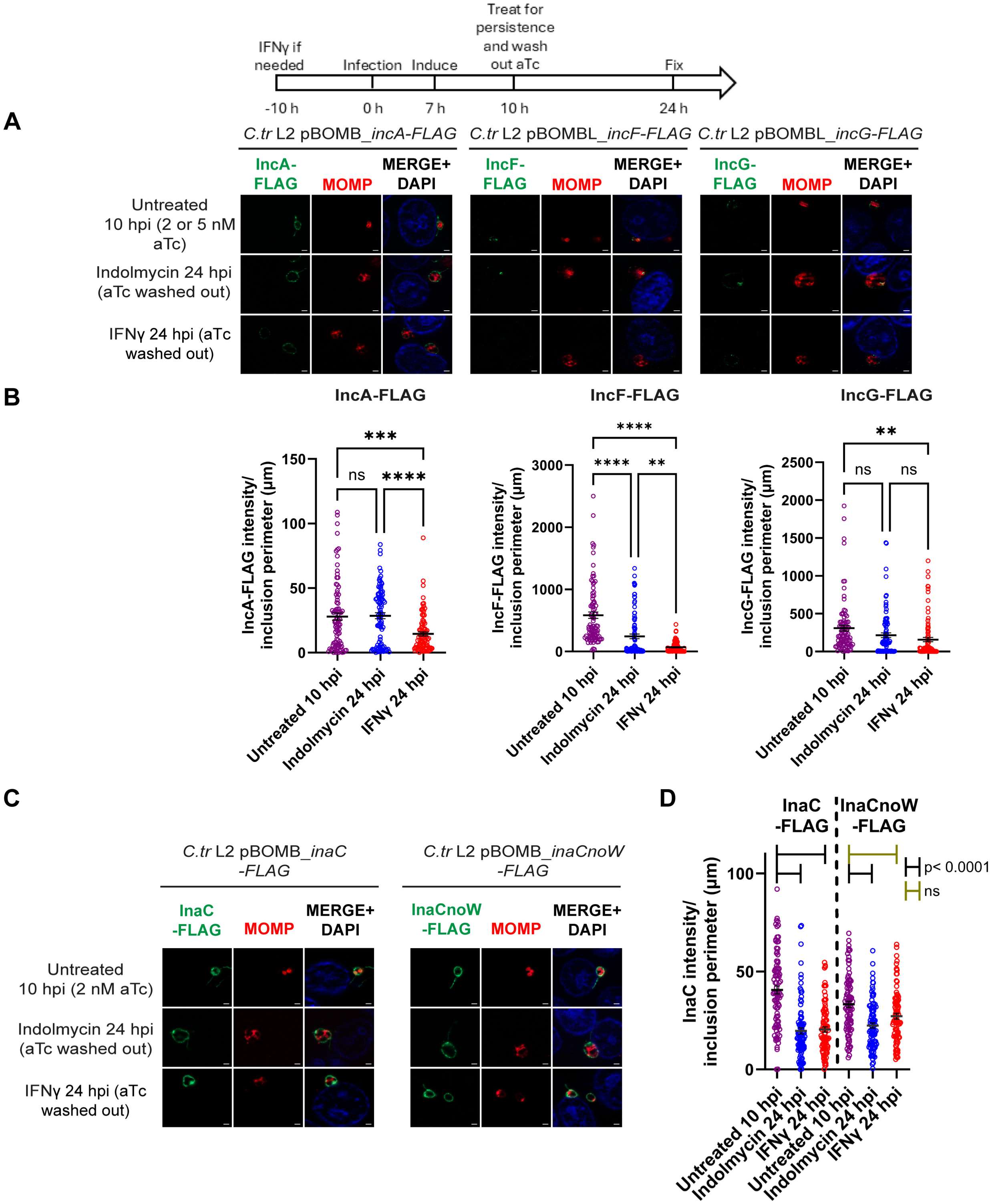
Stability of FLAG tagged Incs is altered during Trp starvation. A diagram representing the experimental design and highlighting important timepoints is shown. At 7 hpi, 2 nM aTc (IncA-FLAG, InaC-FLAG, and InaCnoW-FLAG) or 5 nM aTc (IncF-FLAG and IncG-FLAG) was added to induce construct expression. **(A, C)** Samples were fixed at 10 hpi or 24 hpi and processed for indirect immunofluorescence to detect FLAG (green), MOMP (red), and DNA (blue). Scale bar = 2 μm. **(B, D)** Immunofluorescent images were quantified using Fiji/ImageJ software to determine the raw integrated density of FLAG tagged protein, normalized to exposure time if necessary (IncA-FLAG, InaC-FLAG, and InaCnoW-FLAG), and divided by inclusion perimeter (µm), and results (mean and SEM) were graphed and statistically analyzed (ordinary one-way ANOVA with Šidák’s multiple comparisons test) using GraphPad Prism. ns, non-significant; **, p<0.01; ***, p<0.001; ****, p<0.0001. In **D**, gold brackets denote ns, non-significant, and black brackets indicate statistical significance: p<0.0001. Data shown are combined from 3 independent experiments (90 inclusions total per condition).

The effect of Trp-codon content on InaC stability at the IM during Trp starvation was also tested by infecting HEp-2 cells with *C. trachomatis* strains carrying inducible wild-type InaC-FLAG or mutant InaCnoW-FLAG as described above. At the indicated timepoints, cells were fixed and processed for indirect immunofluorescence (Fig. 6C), and intensity of Inc-FLAG constructs on the IM was quantified (Fig. 6D). Also as above, persistence was confirmed by visualization of enlarged organisms (Fig. 6C) within quantifiably small inclusions (Fig. S4). Expression of both InaC-FLAG and InaCnoW-FLAG levels at the IM were similar at 10 hpi, but there was a lower amount of InaCnoW-FLAG compared to the wild-type InaC-FLAG (p=0.0152) (Fig. 6C and D). During persistence, IM-associated levels of both InaC-FLAG and InaCnoW-FLAG were decreased compared to the 10 hpi controls. InaC-FLAG was significantly decreased under both persistence conditions (p<0.0001). InaCnoW-FLAG was significantly decreased during indolmycin-induced persistence (p<0.0001), and while InaCnoW-FLAG was also decreased during IFNγ-induced persistence, this was not statistically significant (p=0.0893). When comparing wild-type InaC-FLAG stability between indolmycin and IFNγ treatments, there was no difference in Inc stability; similarly, mutant InaCnoW-FLAG also showed no difference in stability between indolmycin and IFNγ conditions. Comparing InaC-FLAG and InaCnoW-FLAG levels during indolmycin-induced persistence, there was no significant difference in Inc stability. However, when comparing InaC-FLAG and InaCnoW-FLAG levels during IFNγ-induced persistence, there was statistically significantly more InaCnoW-FLAG on the IM (p=0.0333). These data suggest that Inc expression on the IM is more complex than just the Trp content of each individual Inc. Additionally, these data indicate that host cell processes may also influence Inc content of the IM.

## Discussion

One host defense mechanism used to combat invading pathogens is to limit the availability of critical nutrients like the essential amino acid, Trp. In humans, this occurs via the production of the cytokine IFNγ, which activates target cells to produce the Trp catabolizing enzyme IDO. For Trp auxotrophic pathogens, like *Chlamydia*, this is particularly problematic, as they must acquire Trp from the host. Trp limitation leads to chlamydial persistence, and we have previously explored how the bacterium responds to this stress (19–23, 54). Here, we were interested in understanding how the inclusion is maintained during chlamydial persistence induced by Trp starvation. To this end, two previously established *in vitro* models that induce persistence via Trp starvation with either IFNγ or the antibiotic, indolmycin, were used to examine specific host-chlamydial interactions and to begin to understand the Inc composition of the IM during persistence (20–23, 54). Data from this study demonstrate that *Chlamydia* is able to maintain its niche during Trp limiting conditions because key interactions with the host cell are maintained. Further, Inc composition of the IM is likely limited to Incs that are sufficient to maintain key *Chlamydia*-host interactions, and the overall Inc composition of the IM during persistence is different than what is observed during normal development. Specifically, Inc localization to the inclusion during persistence is dependent on multiple factors, including the Trp content of the individual Inc, the ability of the Inc to be T3Sed, and the stability of the Inc once it localizes to the IM.

For the first time, persistent *Chlamydia* were shown to acquire SM and cholesterol, which correlated with the maintenance of Rab6 trafficking to the IM (Fig. 1). CERT, which has been associated with ER-IM contact sites (55, 56), was also demonstrated to localize to the inclusion during persistence (Fig. 3E and F) and correlated with IncD expression during persistence (Fig. 3A and B). Components of the recycling endosomal pathway, including Rab11-FIP1 and Tf, also localized to the inclusion during persistence (Fig. 2). Combined, these data indicate that during persistence induced by Trp starvation, the inclusion maintains interactions with the Golgi (Fig.1), early interactions with the ER via CERT recruitment (Fig. 3), and the endosomal recycling pathways (Fig. 2). All these pathways have been linked to chlamydial viability.

Chlamydial Incs function both to recruit host factors to the inclusion and stabilize the IM to support chlamydial development. We prioritized examination of Incs for which we had the greatest resources, including antibodies to endogenous proteins. Among our list of tested Inc proteins in *Chlamydia*, InaC is the only Inc enriched in Trp codons with a Trp motif (WW), three other Incs have a low Trp content (IncD, IncF, and CT223), and three other Incs lack Trp codons entirely (IncE, IncG, and IncA) (Table 1). The intensity of the endogenous Incs, IncD, IncE, and IncG, at the IM was either unaffected by Trp starvation or was increased relative to the 11.5 hpi untreated condition, which is consistent with their lack of Trp codons (Figs. 3 and 4). Interestingly, the amount of SNX6 recruited to the IM was unaffected by the increased intensity of IncE on the IM during persistence conditions and at 24 hpi in the unteated condition relative to the 11.5 hpi untreated condition; in addition, the intensity of IncD on the IM negatively correlated with the level of its binding partner, CERT, recruited to the IM (Fig. 3). Given that IncD and IncE are expressed on the IM during persistence, it is likely that SNX6 and CERT are trafficking to the IM as they would during a normal chlamydial infection and that the differences seen here are not biologically significant. Thus, *Chlamydia* can recruit these host cell proteins to the IM during Trp limitation, and presumably, Inc binding and function at the IM is maintained. Meanwhile, Trp starvation decreased detection of endogenous IncA (0 Trp) and InaC (6 Trp) on the IM (Fig. 4). Inducing expression of InaC-FLAG from *C. trachomatis* during persistence resulted in a lack of InaC-FLAG on the IM, or within organisms, similarly to endogenous InaC (Figs. 4 and 5C and D). Intriguingly, mutating all the Trp codons to Phe codons in InaC-FLAG, creating InaCnoW-FLAG, resulted in the expression and localization of InaCnoW-FLAG to the IM during persistence (Fig. 5C and D). These data support the hypothesis that the Trp codon content of Incs can directly impact their expression and localization to the IM during Trp-starvation induced persistence.

In contrast, some of the data suggest that additional factors other than Trp content of the individual Inc are involved in Inc localization to the IM during persistence. These additional factors may include, T3S functionality, stability of the Inc, and temporal expression during the developmental cycle. Concerning temporal expression, induction of persistence at 10 hpi may favor detection of early Incs (e.g., IncD, IncE, and IncG (current study; Figs. 3 & 4) and CT229 (24)) on the IM, as these Incs are expressed before the induction of persistence. Similarly, endogenous IncA (0 Trp) was not detected on the IM during persistence but under normal culture conditions, is expressed 8-10 hours after persistence was induced in these experiments (Fig. 4). However, when IncA-FLAG was induced for expression from *C. trachomatis* 14 hours after the induction of persistence, IncA-FLAG was found on the IM and associated with fibers extending from the inclusion (Fig. 5A and B). These data suggest that chlamydiae are competent to secrete IncA during persistence, which suggests transcriptional control of *incA* expression during persistence. Of note, a previous study found limited changes to *incA* transcript levels during IFNγ-induced persistence when comparing transcript levels between 24-hpi IFNγ-treated samples and 10 hpi untreated samples (1.16 fold change, p=0.890), while there was a 4.97 fold increase in *incA* transcripts in 24 hpi indolmycin-treated samples (p=0.144) (22). Hence, the reason for the lack of IncA on the IM during persistence is not likely linked to its transcript level. Interestingly, induction of expression of either IncF-FLAG or IncG-FLAG from *C. trachomatis* and under the same conditions as IncA-FLAG, did not result in either construct localizing to the IM (Fig. 5A and B). These results differed from the results obtained with endogenous proteins. In those experiments, induction of persistence occurred after the onset of transcription of *incF* and *incG*, but before the onset of transcription of *incA*. Importantly, the IncA-FLAG data indicate that the T3S components required for IncA-FLAG secretion were expressed and functional during indolmycin or IFNγ-induced persistence, indicating that their absence is not the mechanism driving lack of endogenous IncA during persistence. IncF has a single Trp, and IncG has no Trp, which contradicts, or at least confounds, the original hypothesis that Trp content is the most important determinant in protein expression during persistence. These data suggest additional factors, such as the ability of proteins to be type III secreted or protein stability, may impact the overall Inc content of the IM.

The data demonstrating that both IncA-FLAG and InaCnoW-FLAG localize to the IM indicate that chlamydial T3S is functional during persistence induced by Trp starvation (Fig. 5). There are 26 proteins that are thought to comprise the chlamydial T3S apparatus (including basal body, export apparatus, needle, and needle tip) and 11 candidate chaperone proteins, which are required for effector secretion (Suppl. Table S1) (reviewed in (53)). The majority of these proteins, including the Mcsc chaperone (thought to assist in Inc secretion (57)) and Slc1 (chaperone for early effectors like TarP (58–60)), are low in Trp, with an average Trp content of only 0.67%, and thus, are predicted to be expressed during Trp starvation. Previous RNAseq analysis of chlamydial transcription during persistence supports that Mcsc and Slc1 are likely transcribed, but there were not complete data associated with all predicted chaperones (22). The apparatus components that contain greater than 1.3% Trp include CdsJ (basal body component; 1.54% Trp), several components of the export apparatus (CdsS [2.13% Trp], CdsT [2.42% Trp], and CdsV [1.49% Trp]), PknD (serine/threonine kinase; 1.39% Trp), and CopB2 (needle translocator; 1.38% Trp); of these components, CopB2 is redundant to CopB (reviewed in (53)). It is unknown why the *C. trachomatis* genome contains some redundant genes encoding components of the T3S apparatus, but one possibility is to exact effector secretion without the layers of regulation that govern the expression and function of T3S systems in other Gram-negative bacteria (53). A previous study found *copB2* and *pknD* to be down-regulated during IFNγ-induced persistence and *copB1* to be down-regulated during indolmycin-induced persistence (22). Future studies will be designed to directly address the expression and function of T3S system components during persistence caused by Trp-starvation and whether these are linked to chlamydial-specific and/or host-specific processes.

During a natural *Chlamydia* infection, the inflammatory cytokine IFNγ is produced by the innate immune response as well as CD4+ T-cells and is thought to have a protective effect against infection (61). IFNγ is also able to trigger persistence, which complicates treatment and clearance of *Chlamydia* (28). IFNγ action creates a Trp-limiting environment both for the host cell and *Chlamydia*. On the contrary, the target of indolmycin is the bacterial tryptophanyl-tRNA synthetase and, consequently, this drug will affect only *Chlamydia* and not the host cell (33). Thus, the use of both methods to induce chlamydial persistence allows the distinction between host- and chlamydial-specific effects on IM composition. Among the host factors included in our study, we observed a significant difference in Rab11-FIP1 detection at the inclusion between IFNγ- and indolmycin-mediated persistence at 24 hpi (Fig. 2C). Persistence induced by indolmycin caused a significant decrease in localization of Rab11-FIP1 to the IM at 24 hpi compared to untreated cells at 11.5 hpi (p<0.0001) while IFNγ-induced persistence did not cause a significant change in Rab11-FIP1 at the IM at 24 hpi compared to untreated cells at 11.5 hpi. The difference in Rab11-FIP1 localization to the IM between the two persistence conditions was greatest at 24 hpi but was not quite statistically significant (p=0.0674). By 48 hpi, both Trp-depleting treatments led to similar amounts of Rab11-FIP1 at the IM. Altogether, this suggests that chlamydial-specific protein synthesis inhibited by indolmycin significantly decreased the ability of Rab11-FIP1 to localize to the inclusion but only at 24 hpi. While Rab11-FIP1 has been demonstrated to localize to the IM by as early as 12 hpi (46), the exact mechanism of its recruitment to the IM in unknown, particularly regarding possible bacterial mechanisms of recruitment.

When examining endogenous Incs on the IM during persistence conditions, there was increased IncD and IncG on the IM during both persistence treatments at 24 hpi compared to the untreated 11.5 hpi condition (Fig. 3B and 4B). However, the increases in endogenous IncD and IncG on the IM were both only significant during IFNγ-mediated persistence (p=0.012 for IncD and p=0.0052 for IncG), suggesting that targeting host cell tryptophan availability could increase expression and localization of these Incs to the IM. This would not be surprising given that IncD is Trp-poor, IncG lacks Trp altogether, and both are expressed early in development, before persistence was induced in our study. Considering that the total amounts of IncD and IncG on the IM are similar between the two persistence conditions and statistically insignificantly different from one another, it is more likely that persistence does not cause a biologically relevant increase in either IncD or IncG on the IM, regardless of the mechanism of inducing persistence. In the case of IncD, while both persistence conditions at 24 hpi resulted in higher amounts of IncD on the IM compared to the untreated 11.5 hpi condition, levels were still lower than the untreated 24 hpi condition.

Unexpectedly, we observed significant differences regarding Inc protein stability during persistence induced by IFNγ or indolmycin. For example, less IncA-FLAG was detected on the IM in IFNγ-treated cultures after aTc washout compared to inclusions where persistence was induced with indolmycin (p<0.0001), where the quantity of IncA-FLAG on the IM was comparable to that of the untreated 10 hpi control (Fig. 6B). Similarly, less IncF-FLAG was found on the IM during both persistence conditions compared to the untreated 10 hpi condition (p<0.0001), with an even greater decrease found in IFNγ-treated cells than indolmycin-treated ones (p=0.0027 comparing indolmycin and IFNγ conditions) (Fig. 6B). IncG-FLAG followed a similar pattern to IncF-FLAG, but the decrease in IncG-FLAG stability was only statistically significant when comparing the IFNγ-treated cells to the untreated 10 hpi cells (p=0.0037) (Fig. 6B). There has yet to be a study that formally measures the stability of Incs that localize to the IM throughout chlamydial development. However, these data suggest that a host-specific process may play a role in Inc turnover at the IM. While we cannot formally exclude that the FLAG tag itself may be targeted by a host factor induced by IFNγ that mediates this effect, this scenario would be surprising given that FLAG epitope tags have been used to understand protein-protein interactions during IFNγ signaling (62–64). Finally, InaC-FLAG and InaCnoW-FLAG were both less stable under both persistence conditions compared to the 10 hpi untreated samples (Fig. 6D). For InaC-FLAG, both indolmycin and IFNγ treatments resulted in significantly less Inc stability (p<0.0001) while InaCnoW-FLAG stability was significantly decreased only when indolmycin induced persistence (p<0.0001). This could be due to an issue with chlamydial protein synthesis; however, this would be unexpected, since InaCnoW-FLAG has no Trp, and the wild-type InaC-FLAG, which has Trp, did not show a difference in stability between persistence conditions. Given the lack of a statistically significant difference between InaCnoW-FLAG levels on the IM between the persistence conditions, a more likely explanation is that InaC shows less stability on the IM during persistence conditions, regardless of the presence of Trp or the mode of inducing persistence.

Together, our data provide evidence that core host-chlamydial interactions are preserved during persistence. These are the first experiments to determine how the inclusion remains stable and *Chlamydia* remain viable, yet quiescent, during persistence. While we cannot perform a comprehensive study due to limited resources targeting endogenous Incs, our data demonstrate that, during persistence, there is a partial remodeling of the IM as Inc abundance in the IM is changed. Overall, these studies highlight the need for future studies to define chlamydial T3S function and expression as well as Inc protein dynamics on the IM during chlamydial development. A better understanding of these chlamydial processes may lead to novel antimicrobials that can target *Chlamydia* during all stages of development, including during persistence.

## Materials and Methods

### Organisms and cell culture

The HEp-2 (persistence assays) and McCoy (transformation of *C. trachomatis* serovar L2 strain 434/Bu) cells were kind gifts of Dr. Harlan Caldwell (NIH/NIAID). Cells were routinely cultured in Dulbecco’s modified Eagle’s medium (DMEM; Invitrogen, Waltham, MA) containing 10% fetal bovine serum (FBS; Hyclone, Logan, UT) and 10 μg/mL gentamicin (Gibco, Waltham, MA) at 37°C, 5% CO_2_. The HEp-2 and McCoy cells were verified to be mycoplasma negative using the LookOut Mycoplasma PCR Detection kit (Sigma, St. Louis, MO). *C. trachomatis* L2/434/Bu was cultured and purified as previously described (65). Eukaryotic cells were infected with *C. trachomatis* L2 EBs or chlamydial transformants in DMEM with 10% FBS and 10 μg/mL gentamicin by centrifuging at 400 *g* at room temperature for 15 minutes. A multiplicity of infection (MOI) of 1 was used, unless otherwise specified.

### Induction of persistence in *C. trachomatis*

IFNγ-conditioned medium (ICM) was prepared by adding 2 ng/mL IFNγ to uninfected HEp-2 cells for approximately 54 h prior to collection and filtration of the medium. For longer storage, ICM was kept at −20°C. Indolmycin (Cayman Chemical, Ann Arbor, MI) was prepared in dimethyl sulfoxide (DMSO; Sigma) and stored at −80°C. IFNγ (Cell Sciences, Canton, MA) was resuspended to 100 μg/mL in 0.1% bovine serum albumin (BSA; Sigma) diluted in water. Aliquots were kept at −80°C and used only once to avoid freeze-thawing. At 10 hours post infection (hpi), cells were washed twice with Hanks’ Balanced Salt Solution (HBSS; Gibco). *To induce persistence using the tryptophanyl-tRNA synthetase inhibitor*, 120 μM indolmycin in ICM was added after the washes with HBSS. *To induce persistence using recombinant human IFNγ*, 10 h prior to infection, 0.5 ng/mL was added to the cells. To induce persistence in *C. trachomatis,* DMEM medium was replaced by the Trp-depleted medium, ICM, after the washes with HBSS at 10 hpi, as previously described (21).

### Transformation of *Chlamydia trachomatis*

Plasmid-free *C. trachomatis* serovar L2 (-pL2) EBs were prepared and used for transformation as previously described (66). Briefly, 10^6^ *C. trachomatis* serovar L2 (-pL2) EBs were incubated with 2 μg of plasmid in a volume of 50 μL Tris-CaCl_2_ buffer (10 mM Tris, 50 mM CaCl_2_, pH 7.4) for 30 min at room temperature. McCoy cells cultured in a 6-well plate were washed with 2 mL HBSS, and 1 mL HBSS was added back into each well. The transformation inoculum, mixed to 1 mL HBSS, was then used to infect a confluent monolayer of McCoy cells by centrifugation at 400 *g* for 15 min at room temperature followed by incubation at 37°C for 15 min. The inoculum was then aspirated, and DMEM containing 10% FBS and 10 μg/mL gentamicin was added to the well. At 8 hpi, the medium was replaced with DMEM containing 1 or 2 U/mL penicillin G and 1 μg/mL cycloheximide. The plate was incubated at 37°C for 48 h; then, the transformants were harvested (T_1_) to infect a new McCoy cell monolayer. Transformants were passaged and subcultured every 48 hpi until a population of penicillin-resistant bacteria was observed. EBs were harvested and frozen in sucrose-phosphate (2SP) (66) solution and stored at −80°C. To verify each transformant, plasmid DNA was isolated from infected cells for subsequent restriction digest and sequencing.

### Cloning

The plasmids and primers used in this study are described in Suppl. Table 2. The *C. trachomatis* L2 pBOMB4-Tet-IncA-FLAG (CTL0374) strain (67), the *C. trachomatis* L2 pBOMB4-Tet-InaC-FLAG (CTL0184) strain (68), and the *C. trachomatis* L2 pBOMBmC-*gfp(ssrA_VDD)* strain (41) were described previously. The *InaCnoW-FLAG*, the *incF-FLAG*, and the *incG-FLAG* constructs for chlamydial transformation were created using the HiFi DNA Assembly Cloning Kit (New England Biolabs, Ipswich, MA). The backbone used was pBOMB4-Tet (pBOMB4-Tet is a kind gift from Ted Hackstadt) (69) and its derivative pBOMBL (70). *InaCnoW-FLAG* construct was created for this study by amplifying mutant *inaC-FLAG* without Trp codons (*inaCnoW*) by PCR with Phusion DNA polymerase using a gBlock fragment containing a FLAG tag (Integrated DNA Technologies, Coralville, IA) as a template. gBlock was designed based on the *C. trachomatis* serovar L2 434/Bu *ctl0184* gene, and *noW-FLAG* mutant was obtained by exchanging the TGG codons with TTT or TTC codons encoding phenylalanine residues. The PCR products were purified using a PCR purification kit (Qiagen, Hilden, Germany). *noW-FLAG* was cloned onto the EagI/KpnI sites of the plasmid pBOMB4-Tet using the HiFi DNA Assembly Cloning Kit (NEB). To make the *incF-FLAG* and *incG-FLAG* constructs, a gBlock fragment including a flexible glycine-glycine-glycine-serine (repeated twice) linker and the FLAG tag was first designed. Then, primers were designed to amplify *incF* (*ctl0372*) or *incG* (*ctl0373*) with overlapping regions to the pBOMBL vector in 5′ and the linker in 3′. Then *incF-FLAG* and *incG-FLAG* were cloned onto the EagI/SalI sites of the plasmid pBOMBL using the HiFi DNA Assembly Cloning Kit. In all cases, the product of the HiFi reaction was transformed into DH10β *E. coli* competent cells (NEB) and plated on LB agar plates containing 100 μg/mL ampicillin and 0.4% glucose. Plasmid was subsequently isolated from individual colonies grown overnight in LB by using a miniprep kit (Qiagen) and confirmed by restriction digest and sequencing.

### Antibodies

Primary antibodies used included the following: mouse anti-CT223 (kind gift from R. Suchland, University of Washington, WA, and D. Rockey, Oregon State University, OR); sheep anti-IncA (made to order by Seramum Diagnostica GmbH, Heidesee, Germany); rabbit anti-InaC, rabbit anti-IncD, rabbit anti-IncE, and rabbit anti-IncG (kind gifts from T. Hackstadt, NIAID, Rocky Mountain Laboratories, Hamilton, MT); goat anti-MOMP (Meridian Life Sciences; B65266G; Memphis, TN) and mouse anti-MOMP (Meridian Life Sciences; C01365M; Memphis, TN); chicken polyclonal anti-CERT (Sigma; GW22128B); mouse anti-Sorting nexin-6 (SNX6) clone D-5 (Santa Cruz Biotechnology; sc-365965); rabbit anti-Rab6 (Cell Signaling, Danvers, MA; 9625s); rabbit anti-Rab11-FIP1 (EMD Millipore/Sigma; HPA035960); mouse anti-FLAG (Millipore/Sigma; F1804). Secondary antibodies used for immunofluorescence were donkey anti-mouse-, rabbit-, sheep, or goat-488, donkey anti-mouse-, goat-, rabbit-, or chicken-594 (Jackson ImmunoResearch, West Grove, PA and Invitrogen).

### Indirect immunofluorescence

HEp-2 cells seeded onto glass coverslips in 24-well plates were infected at an MOI of 1 with *C. trachomatis* serovar L2 wild-type or *C. trachomatis* serovar L2 with pBOMBmC-GFP(VAA) transformed into *C. trachomatis* L2 -pL2 (41) (SM trafficking only). Cells were fixed in 100% methanol for 5 min (endogenous Incs and Inc_FLAG strain samples) or were fixed in 4% paraformaldehyde in 1x Dulbecco’s phosphate-buffered saline (DPBS) for 15 min at room temperature to study host markers. Cells fixed with 4% paraformaldehyde were permeabilized with 0.1% Triton X-100 in 1x DPBS for 5 min at room temperature. Coverslips were then processed using indirect immunofluorescence with the appropriate primary and secondary antibodies (see list above). DAPI was used to stain DNA. Primary antibodies were stained overnight at 4°C or for 1 h at 37°C. All secondary antibodies were stained for 1 h at 37°C. Coverslips were washed two times with 1x DPBS between each antibody incubation step. Coverslips were mounted using ProLong Gold antifade mounting medium (Life Technologies, Carlsbad, CA). Images were acquired at 100x magnification using a Zeiss AxioImager Z2 microscope with or without ApoTome.2 as indicated in figure legends and using identical exposure times between conditions (except SM trafficking). Images were compiled in Adobe Photoshop.

### Visualization of SNX6, CERT, Rab6, and Rab11-FIP1 during persistence mediated by indolmycin+ICM or IFNγ

To visualize Sorting-nexin 6 (SNX6) localization, coverslips were methanol fixed at 11.5 hpi and 24 hpi and stained for immunofluorescence to observe expression of SNX6 (red), MOMP (green), or DNA (DAPI; blue). To visualize ceramide transfer protein (CERT) localization, coverslips were fixed with 4% paraformaldehyde at 24 hpi, permeabilized with 0.1% Triton X-100, and then stained for immunofluorescence to observe expression of CERT (red), MOMP (green), or DNA (DAPI; blue). To visualize Rab6 and Rab11-FIP1 localization, coverslips were fixed with 4% paraformaldehyde at 24 hpi and 48 hpi, permeabilized with 0.1% Triton X-100, and then stained for indirect immunofluorescence to observe expression of Rab6 (red), Rab11-FIP1 (red), MOMP (green), or DNA (DAPI; blue).

### NBD-ceramide labeling during persistence mediated by indolmycin+ICM or IFNγ

At 4 hpi and 22.5 hpi, cells were labeled with NBD-ceramide for 30 minutes, essentially as previously described (34, 35). Labeled cells were incubated for 6 h (4 hpi NBD-ceramide treatment), or 4 h (22.5 hpi NBD-ceramide treatment) in medium containing FBS (back-exchange medium) to remove fluorescent lipid that had not been incorporated into chlamydial inclusions. Cells were imaged live using brightfield and the 488 and 594 fluorescent channels on the Zeiss Axio Imager.Z2 without the ApoTome.2 with x63 magnification. Fiji/ImageJ was used to measure intensity of NBD-sphingomyelin and inclusion area for 25 inclusions per condition for each biological replicate. Raw integrated density measurements were normalized to exposure time (in milliseconds), then divided by the area of the inclusion (μm^2^) to account for inclusion size.

### Cholesterol labelling during persistence mediated by indolmycin+ICM or IFNγ

To visualize intracellular cholesterol, coverslips were fixed with 4% paraformaldehyde at 11.5 hpi and 24 hpi, and fixed cells were incubated with 50 μg/mL filipin III (Cayman Chemical) in DPBS + 1% BSA for 1 h at room temperature as previously described (71). Cells were then processed using indirect immunofluorescence to observe SNX6 (green) and MOMP (red) expression. Coverslips were imaged using Zeiss Axio Imager.Z2 without ApoTome.2 with x100 magnification.

### Transferrin labeling during persistence mediated by indolmycin+ICM or IFNγ

To study transferrin-positive vesicles, 24-well plates of HEp-2 cells were infected at an MOI of 1 with *C. trachomatis* serovar L2. Aberrancy was induced or not at 10 hpi as described above. At 11.5 hpi and 24 hpi, cells were labeled as previously described (46). Briefly, HEp-2 cells were treated with 25 μg/mL human transferrin conjugated to a 488 fluorophore (Life Technologies/Thermo Fisher) added directly to culture medium. Cells were incubated with transferrin-488, culture media was aspirated, and cells were washed once with 1x DPBS prior to fixation. Samples were then processed for indirect immunofluorescence essentially as described above.

### Visualization of endogenous Incs during aberrancy mediated by indolmycin+ICM or IFNγ

Aberrancy was induced or not with 120 μM indolmycin or IFNγ following the procedures described above. At 11.5 hpi, 16 hpi (for InaC samples only), or 24 hpi, coverslips were fixed. Bacteria were detected using goat anti-MOMP or mouse anti-MOMP antibodies (Meridian). Endogenous Incs were detected using primary antibodies and secondary antibodies described above. All images for a single Inc were captured using the same exposure time (milliseconds) across all biological replicates, unless otherwise stated. Fiji/ImageJ software (72) was used to quantify Incs fluorescence intensity. Thirty inclusions per condition for each replicate were measured. Regions of interest (ROI) were defined in merged fluorescent images, and the raw integrated density was measured from the defined ROI in the images of the individual channels: CT223-488, InaC-488, IncA-488, IncD-488, IncE-488, and IncG-488 channels. In total, the raw integrated density was measured for 90 inclusions and normalized to the perimeter of the inclusion (µm) and expressed as Inc intensity/inclusion perimeter (µm). Data were plotted, and statistically analyzed by one-way ANOVA test with Šidák’s multiple comparisons test, as appropriate, using GraphPad Prism software.

### Analysis of construct secretion or stability during aberrancy mediated by tryptophan starvation

HEp-2 cells seeded on glass coverslips in 24-well plates were infected at an MOI of 1 with *C. trachomatis* L2 pBOMB-*incA*-FLAG, pBOMBL-*incF*-FLAG, pBOMBL-*incG*-FLAG, pBOMB-*inaC*-FLAG, or pBOMB-*inaCnoW*_FLAG strain. Aberrancy was induced or not with 120 μM indolmycin or IFNγ following the procedures described above. *To study construct secretion*, induction of IncA-FLAG, IncF-FLAG, IncG-FLAG, InaC-FLAG, and InaCnoW-FLAG expression was performed by the addition of 2 nM aTc at 24 hpi and allowed to proceed for 6 h. At 30 hpi, coverslips were fixed in 100% methanol. *To study construct stability*, induction of IncA-FLAG, InaC-FLAG, and InaCnoW-FLAG expression was performed by the addition of 2 nM aTc at 7 hpi, and induction of IncF-FLAG and IncG-FLAG expression was performed by the addition of 5 nM aTc at 7 hpi. At 24 hpi, coverslips were fixed in 100% methanol. All fixed cells were processed for indirect immunofluorescence and imaged at the same exposure time across all replicates. Fiji/ImageJ software was used to quantify FLAG fluorescence intensity. Thirty inclusions per condition for each replicate were measured. Regions of interest (ROI) around the inclusion membrane were defined in merged fluorescent images. The raw integrated density from the defined ROI in the images of the FLAG-488 channel was measured. In total, the raw integrated density was measured for 90 inclusions and normalized to the perimeter of the inclusion (µm) and expressed as FLAG/inclusion perimeter (µm). Data were plotted, and statistically analyzed by ordinary one-way ANOVA test with Šidák’s multiple comparisons using GraphPad Prism software.

## Supplemental material

Supplementary data are available online.

### AUTHOR CONTRIBUTIONS

C.R., E.A.R., and S.P.O. designed and performed the experiments. E.A.R. and S.P.O. supervised research. The data were analyzed by C.R., R.E.W, E.A.R., and S.P.O. C.R. and R.E.W. prepared the figures. C.R. wrote the manuscript, which was edited by E.A.R., R.E.W., and S.P.O.

### FUNDING

This work was supported by funding from the U.S. National Institutes of Health grant 1R01AI132406 to E.A.R. and S.P.O.

## Acknowledgements

The authors would like to acknowledge H. Caldwell (NIH) for eukaryotic cell lines. We thank R. Suchland (University of Washington, WA) and D. Rockey (Oregon State University, OR) for the anti-CT223 antibody. We thank T. Hackstadt (RML/NIAID) for providing the pBOMB4-Tet plasmid and the anti-IncD, the anti-IncE, the anti-IncG, and the anti-InaC antibodies. We thank Lindsey Knight for technical assistance. The authors are thankful to the members of the Rucks/Ouellette lab for helpful comments and suggestions regarding this work.

## Supplemental Material

**Supplementary Table 1. Tryptophan content of the T3S apparatus and chaperones in** *Chlamydia trachomatis* **L2.**

**Supplementary Table 2. List of Plasmids, Strains, and Primers used in this study.**

**Supplementary Figure 1. Diagrams and Inclusion Sizes related to Golgi trafficking experiments. (A, C)** A diagram that depicts the experimental design and includes important timepoints within the experiments. **(B)** The inclusion area (μm^2^) of the NBD-ceramide-labeled inclusions was quantified using Fiji/ImageJ software. The results were graphed and statistically analyzed using GraphPad Prism. Horizontal lines indicate means, and vertical lines indicate standard errors of the means. Statistical significance was determined by ordinary one-way ANOVA with Šidák’s multiple comparisons. **, p<0.01; ****, p<0.0001. **(D)** The inclusion perimeter (μm) was determined, and results were graphed and statistically analyzed as in **B**. Horizontal lines indicate means, and vertical lines indicate standard errors of the means. Statistical significance was determined by ordinary one-way ANOVA with Šidák’s multiple comparisons. ns, non-significant; ****, p<0.0001.

**Supplementary Figure 2. Localization of the CT223, IncG, and InaC Incs at the inclusion membrane during normal developmental cycle.** HEp-2 cells were infected with *C. trachomatis* serovar L2 at an MOI of 1. Samples were fixed at 24 hpi and processed for indirect immunofluorescence to detect Incs (green), MOMP (red), and DNA (blue). Scale bar = 2 μm.

**Supplementary** Figure 3**. Persistence mediated by Trp-starvation negatively impacts inclusion growth compared to untreated control during Inc secretion experiments. (A, B)** Inclusion perimeters (μm) were quantified using Fiji/ImageJ software, and results were graphed and statistically analyzed using GraphPad Prism. Horizontal lines indicate means, and vertical lines indicate standard errors of the means. Statistical significance was determined by ordinary one-way ANOVA with Šidák’s multiple comparisons. ns, non-significant; *, p<0.05; ****, p<0.0001.

**Supplementary Figure 4. Persistence mediated by Trp-starvation negatively impacts inclusion growth compared to untreated control during Inc stability experiments. (A, B)** Inclusion perimeters (μm) were quantified using Fiji/ImageJ software, and results were graphed and statistically analyzed using GraphPad **P**rism. Horizontal lines indicate means, and vertical lines indicate standard errors of the means. Statistical significance was determined by ordinary one-way ANOVA with Šidák’s multiple comparisons. **, p<0.01; ****, p<0.0001.

